# Human Nerve-on-a-Chip. Engineering 3D functional human peripheral nerve *in vitro*

**DOI:** 10.1101/458463

**Authors:** Anup D. Sharma, Laurie McCoy, Elizabeth Jacobs, Hannah Willey, Jordan Q. Behn, Hieu Ngyuen, Brad Bolon, J. Lowry Curley, Michael J. Moore

## Abstract

Development of “organ-on-a-chip” systems in the space of the nervous system is lagging because of lack of availability of human neuronal & glial cells and the structural complexity of the nervous system. Furthermore, a sizeable challenge in the drug development pipeline and an often-cited reason for failure in a vast number of clinical trials conducted for neuropathological ailments is the lack of translation from animal models. A preclinical *in vitro* model capable of mimicking the function and structure of the human nervous system is well desired. We present an *in vitro* biomimetic model of all-human peripheral nerve tissue capable of showing a robust neurite outgrowth (~5mm), myelination of human induced pluripotent stem cells (iPSCs)-derived neurons by primary human Schwann cells and evaluation of nerve conduction velocity (NCV), previously unrealized for any human cell-based *in vitro* system. This Human Nerve-on-a-chip (HNoaC) system is the first microphysiological system of human peripheral nerve which can be used for evaluating electrophysiological and histological metrics, which are gold-standard assessment techniques previously only possible with *in vivo* studies.

## Introduction

The last decade has observed a rapid pace in development of *in vitro* microphysiological systems, including organs-on-chips, for mimicking human tissue physiology and, thus, improving the predictivity of preclinical drug screens ^1,2^ and reducing the high rate of late-stage clinical trial failure ^3^. Due to the complexity of the nervous system and the neuron-glia interplay, *in vitro* neural microphysiological systems to evaluate potential human responses to novel therapeutic candidates lag those of other organs. Peripheral nerves in particular lack appropriate human-relevant *in vitro* models. Thus far, animal testing remains the gold standard as they are the only models capable of supporting the standard clinically relevant metrics used for assessing peripheral neurotoxicity, namely electrophysiological and histopathological data ^4^.

In this study, we describe an *in vitro*, microengineered, biomimetic, all-human peripheral nerve (Human-Nerve-on-a-Chip [HNoaC]) comprised of induced pluripotent stem cell (iPSC)-derived neurons (hNs) and primary human Schwann cells (hSCs) that can provide data suitable for integrated nerve conduction velocity (NCV) and histopathological assessments. This all-human system is an significant extension of our *in vitro* “Nerve-on-a-chip” (NoaC) platform previously developed using embryonic rat dorsal root ganglion (DRG) neurons and rat SCs ^5^. To our knowledge, this combination of hNs and hSCs has not previously been achieved for any other stem cell-based *in vitro* neural system. This model mimicked robust axonal outgrowth (~5mm), showed first evidence ever of human Schwann cell myelination of human iPSC-derived neurons and first ever proof of nerve conduction velocity testing in an all human *in vitro* system, like *in vivo* models. Therefore, the innovative HNoaC model of human peripheral nerve has the potential to accelerate the field of human disease modeling, drug discovery, and toxicity screening.

## Materials and Methods

### Schwann cell culture

A T-75 culture flask (353136; Corning, Corning, NY) was prepared by coating with a sterile-filtered, 0.1% poly-L-ornithine (PLO; Sigma-Aldrich, St. Louis, MO) solution in sterile water (Sigma-Aldrich, St. Louis, MO). The flask was then washed four times with sterile water. 7.5 mL of a 10μg/mL Laminin (Sigma-Aldrich, St. Louis, MO) in phosphate-buffered saline (PBS; Caisson Labs, Smithfield, UT) was added to the flask, which was held at 4°C overnight. The Laminin solution was aspirated, and 15 mL of culture medium was directly placed into the T-75 culture flask, which was then equilibrated in a 37°C incubator before cell plating.

Human Schwann Cells (hSC) medium was purchased from ScienCell (Carlsbad, CA). The human Schwann cell line (cat. No. 1700; ScienCell) was received in a cryovial with reportedly more than 5 x 10^5^ cells/mL. The vial was removed from cryopreservation and thawed in a 37°C water bath. The contents of the vial were dispensed evenly onto the PLO/Laminin-coated T-75 Flask. The culture was left undisturbed at 37°C in a 5% CO_2_ atmosphere for at least 16 hrs to promote attachment and proliferation. Culture medium was changed every 24 hours. Upon reaching 80% confluency, the hSCs were passaged by using 3 mL of Accutase^®^ (Sigma-Aldrich), which was added to the flask for 3 mins at 37°C. Once cells detached completely, 8 mL of hSC medium was placed in the flask. The 11 mL solution of detached hSCs was moved to a 15 mL conical tube and spun at 200 x g (Eppendorf 5810 R centrifuge, 18cm radius; Eppendorf, Hamburg, Germany) for 5 minutes at room temperature (RT, approximately 22°C). The supernatant was aspirated, and the pellet was resuspended in 1 mL of hSC culture medium. The cells were counted using a conventional hemocytometer (Hausser Scientific, Horsham, PA).

### Motor neuron culture

iCell^®^ Motor Neurons (hNs) medium was prepared using 100 mL of iCell^®^ Neurons Base Medium (FUJIFILM Cellular Dynamics, Inc, Madison, WI) supplemented with 2 mL of iCell^®^ Neural Supplement A (FUJIFILM Cellular Dynamics, Inc) and 1 mL of iCell^®^ Nervous System Supplement (FUJIFILM Cellular Dynamics, Inc). To prepare for thawing motor neurons, hNs medium was warmed to RT and 1 mL of hNs medium was added to a sterile 50 mL conical tube. One vial of iCell^®^ Human Motor Neurons (hNs; FUJIFILM Cellular Dynamics, Inc) was thawed in a 37°C water bath for approximately 2 mins and 30 seconds. The vial contents were transferred to the 50 mL conical tube containing 1 mL of hNs medium, drop-wise with a swirling motion, to mix the cell solution completely and minimize osmotic shock on thawed cells. The cell vial was then rinsed with 1 mL of hNs medium and transferred to the 50 mL tube. The volume of the solution was then brought to 10 mL by slowly adding hNs medium to the 50 mL centrifuge tube dropwise (2-3 drops/sec) while swirling. The cell solution was then transferred to a 15 mL conical tube and centrifuged at 200 x g for 5 mins at RT. Supernatant was aspirated, and cells were resuspended in 1 mL of hNs medium by flicking the tube and then pipetting up and down 2-3 times. A 10 μL sample of cell solution then was taken to perform a cell count using a hemocytometer.

### Spheroid Fabrication

A non-treated, clear, “U” round bottom, 96 well, spheroid microplate (4515; Corning) was used for both monocultures of human Neurons (hNs) and human Schwann cells (hSCs) as well as co-cultures of hNs/hSCs. The concentration in cells/μL of media was calculated for both hSCs and hNs to permit calculation of the volumes needed to produce spheroids of the following sizes and compositions: hNs mono-cultures - 100000, 75000, 50000, or 25000; hSCs mono-cultures - 75000, 50000, or 25000; and co-cultures, 75000 hNs with either 75000, 50000 or 25000 hSCs.

The volume calculated was added to microwell plates, then the volume of each well was brought to 200 μL by adding media warmed to 37°C. The spheroid microplate was then centrifuged at 200 x g for 5 mins and placed in a 37°C incubator in a 5% CO_2_ atmosphere until spheroid formation was observed (typically around 48 h). hNs medium was replenished every other day, at a half changing of 95 μL, and replacing with 100 μL of fresh, warmed (to 37°C) hNs medium.

### Fabrication of 3D dual-hydrogel nerve growth constructs

A dual-hydrogel scaffold was created on the membranes of Transwell^®^ inserts (0.4μm/PES; Corning) using a micro-photolithography technique similar to methods previously described^6^. All solutions were created with sterile-filtered PBS unless otherwise noted. The outer cell-restrictive (i.e., growth-resistant) photo-translinkable hydrogel was created using a solution of polyethylene glycol dimethacrylate 1000 (PEGDMA; Polysciences, Warrington, PA) and lithium phenyl-2,4,6-trimethylbenzoylphosphinate (LAP; Sigma Aldrich). First, 10% w/v PEGDMA and 1.1 mM LAP solutions were created and mixed in a 1:1 solution. The resulting solution was sterile-filtered and added to Transwell^®^ inserts placed in a volume of 0.6 mL while positioned under the lens of a Digital Micromirror Device (DMD, PRO4500 Wintech Production Ready Optical Engine; Wintech Digital Systems Technology Corp, Carlsbad, CA) on Rain-X (ITW Global Brands, Glenview, IL)-treated glass slides (Fig. 1). The mask and polymerization parameters were selected using commercial software (DLP Lightcrafter 4500 Control Software, Texas Instruments, Dallas, TX), and irradiation of the photo-translinkable solution occurred for 28-32 seconds using the ultraviolet light of 385nm wavelength. After treatment, excess PEGDMA/LAP solution was removed from the insert and from within the void created by the photomask. The constructs were then washed using 2% Antibiotic/Antimycotic wash buffer (Thermo Fischer Scientific, Walton, MA) three times on the top and bottom of the insert for 10 minutes each. Wash buffer was removed from the insert and the inner keyhole-shaped channels. The void was carefully filled with 8% Growth Factor-Reduced Matrigel^®^ Matrix (Corning) to create a cell-permeable scaffold and allowed to polymerize in a 37°C incubator.

**Figure 1.**
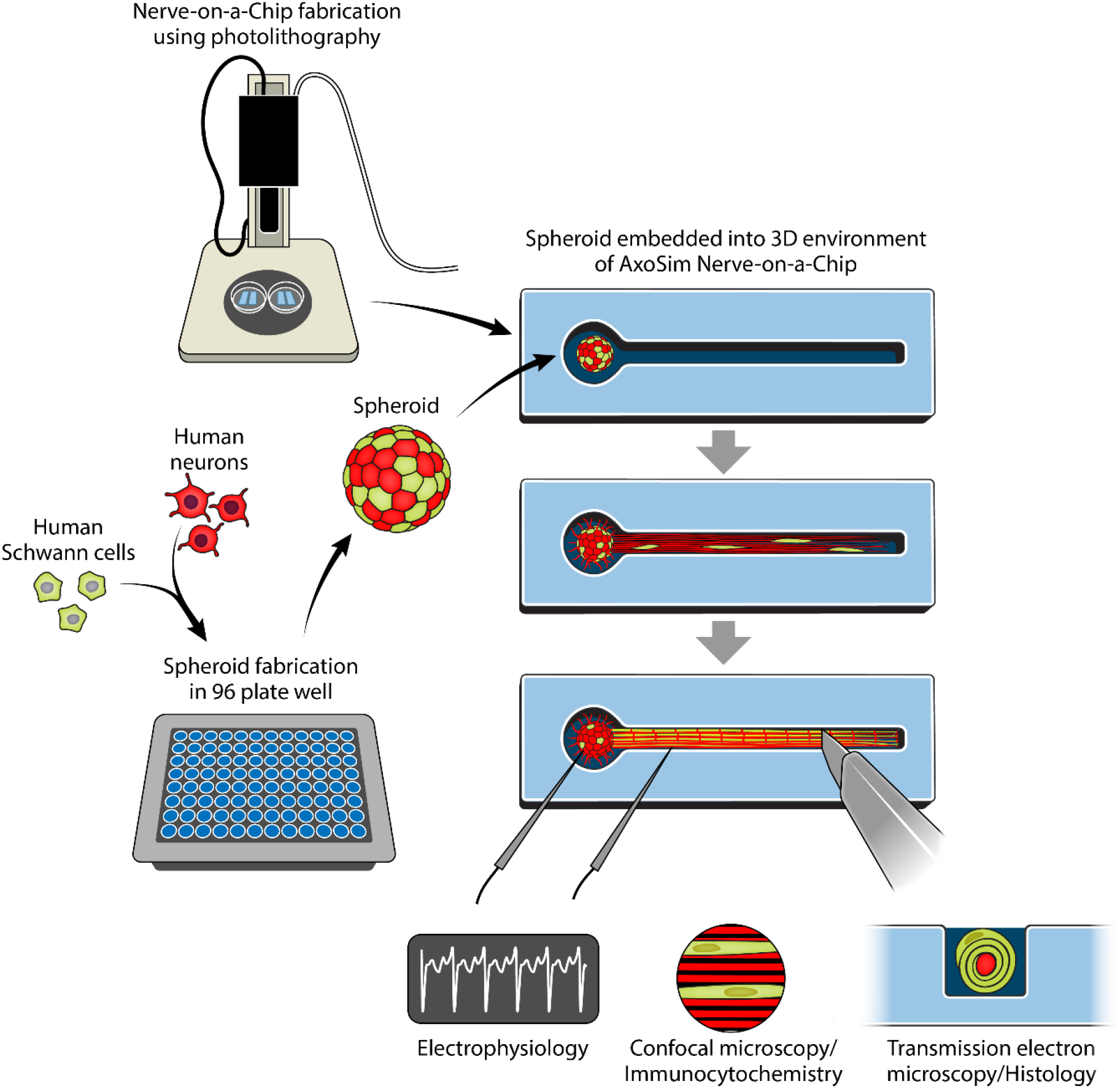
Study design showing the process of fabricating a Human Nerve-on-a-Chip (HNoaC) along with various means by which the system may be characterized (Graphic illustration by Anita Impagliazzo).

### Transferring spheroids to hydrogel construct

Two types of media were created using hNs medium (described above) to induce myelination in 3D constructs. A Pre-myelination medium was created using hNs medium, 10% of HyClone Characterized Fetal Bovine Serum (FBS; LaCell LLC, New Orleans, LA), and 1% Antibiotic-Antimycotic buffer. A Myelination medium was created with hNs medium, 10% FBS, 10ng/mL of recombinant rat beta-Nerve Growth Factor (NGF; R&D Systems, Minneapolis, MN), and 50μg/mL of L-ascorbic acid (Sigma-Aldrich). After formation, spheroids were transferred from the microplate using a pipette and placed onto a 35mm tissue culture-treated dish (Cell Treat, Pepperell, MA) in a droplet of hNs medium. Spheroids were then placed into the 3D construct “bulb portion” (Fig. 1) within the Matrigel, using sterilized Dumont #5 fine-tipped forceps (11295- 10; Dumont, Montignez, Switzerland). 1.5 mL of Pre-myelination medium was finally placed under the Transwell^®^ membrane of the 6-well plates, and the loaded hydrogel constructs were placed in a 37°C incubator in a 5% C02 atmosphere for culture. Half-changes of the medium were performed every other day. The constructs were kept in Pre-myelination medium for 1 week before being switched to Myelination medium for 3 weeks.

### Immunocytochemistry

All wells in the 6-well culture plate were fixed with 4% paraformaldehyde (PFA; Electron Microscopy Sciences, Hatfield, PA), pH 7.4, for 30 min at RT and then washed with PBS 4 times for 15 minutes each. Fixed samples were then placed in a 1X blocking solution containing PBS; 5% normal goat serum (Jackson ImmunoResearch, West Grove, PA); 0.2% Triton-X-100 (Sigma-Aldrich); and 0.4% bovine serum albumin (Sigma-Aldrich) for one hour at RT, followed by labeling with the following primary antibodies overnight in blocking solution at 4°C: rabbit-α- s100 (ab868, 1:400; Abcam, Cambridge, MA); or mouse-α-βIII Tubulin (ab78078, 1:500; Abcam). Rabbit-α-myelin basic protein (MBP, ab133620, 1:500; Abcam) was also used in a separate trial under the same incubation conditions.

The following day, wells were washed with PBS 4 times for 8 min each at RT. The plate was then labeled with secondary antibodies, Alexa 488 goat anti-rabbit IgG (1:300, Abcam) or Alexa 568 goat anti-mouse IgG (1:300, Abcam), and DAPI (1:200, Sigma-Aldrich). Secondary antibodies and DAPI was dissolved in 1X blocker solution for 90 minutes in the dark at RT. The plate was washed 5 times, for 8 mins each, with PBS in the dark at RT. The plate was then parafilmed, foiled, and kept at 4°C until microscopy was performed using a Nikon A1 confocal microscope (Nikon, Tokyo, Japan).

### Plastic resin embedding

All materials used for embedding were purchased from Electron Microscopy Sciences unless otherwise noted and were handled under a chemical flow hood and used with recommended personal protective equipment. Hydrogel constructs were removed from culture and washed three times on both sides of the transmembrane well with PBS at RT prior to fixation. The hydrogel constructs were then soaked in a solution of 4% PFA/0.5% glutaraldehyde for 30 minutes at RT. Secondary fixation and staining of cellular lipids was achieved by post-fixation with 1% osmium tetroxide in PBS, pH 7.4, for 2 hours under dark conditions at RT. The constructs were then washed with PBS 3 times for 15 minutes prior to counterstaining with 2% aqueous uranyl acetate for 30 minutes under dark conditions at RT. Dehydration was done with graded ethanol washes at RT, beginning with a 10-minute wash with 50% ethanol/PBS, a 10-minute wash with 70% ethanol/PBS and an overnight wash with 90% ethanol/PBS. The following day, the constructs were washed twice with 100% ethanol for 30 minutes at RT.

With a scalpel, hydrogel constructs were dissected individually from the transmembrane wells, without removal of the PEGDMA, under a dissecting microscope. Constructs were placed in Flat Embedding Molds (EMS 70902, Electron Microscopy Sciences). Remaining ethanol was given time to evaporate from the fixed hydrogels before replacement with infiltration medium consisting of a 1:1 mixture of Spurr’s resin (Low Viscosity Embedding Media Spurr’s Kit; Electron Microscopy Sciences) and propylene oxide. Infiltration medium was left for 75 minutes before it was replaced by 100% Spurr’s Resin, which was cured overnight in a 70°C oven and for 48 more hours at RT before ultramicrotome sectioning.

### Sectioning and transmission electron microscopy (TEM)

Sectioning and TEM evaluation were performed at the Shared Instrumentation Facility (SIF) at Louisiana State University (Baton Rouge, LA). Ultrathin sections were cut to a thickness of 80- 100 nm at four locations within the HNoaC specimen: within the bulb of the tissue, where the bulb met the channel and the proximal channel (i.e., near the bulb), and distal channel (Fig. 1). Sections were placed on Formvar carbon-coated copper grids, 200 mesh, and impregnated with metal by floating on droplets of 2% uranyl acetate for 20 mins at RT. They were then rinsed with deionized water droplets 3 times, for 1 min. To visualize, a JEOL 1400 TEM (Peabody, MA) was used with an accelerating voltage of 120 kV at varying magnifications.

### Histomorphometric analysis

Metrics acquired from TEM images of HNoaC cross sections included axon diameter and G-ratio (i.e., the ratio of the axon diameter to the diameter of the whole fiber [axon + myelin sheath]). Axon diameter and G-ratio were elucidated by two different, independent, blinded researchers by measuring both unmyelinated axons and axons encircled by 3 or more layers of dark myelin wrapping. G-ratio and axon diameter were measured using the scale, threshold and measure functions in Fiji ^7^. The G-ratio metrics were calculated by randomly sampling 10 images to find axons with 3 or more myelin laminae, while the unmyelinated fibers were measured by randomly sampling 10 images of axons from the distal channel. Axon diameter was measured by using a thresholding function to find the total area of the axon. The diameter was then calculated from area by assuming the axon was circular. G-ratio calculations were based on a simple linear estimation of inner axon diameter while the outer diameter of the whole fiber (consisting of the axon and surrounding dark-stained myelin lamellae) was calculated by taking an average of the smallest and largest diameter of a given nerve fiber. This averaging technique to obtain the outer diameter was needed because the proximity of the myelin layers was inconsistent along the entire circumference of the myelin sheath. G-ratio was calculated by taking the inner diameter over the average outer diameter. The large nucleated bodies of Schwann cells were excluded when measuring the outer extent of the myelin sheath.

### Electrophysiology

After a month in co-culture, the Transwell^®^ insert with reconstituted nerve was placed on a stage for electrophysiological testing. Two tubes, one for dispensing and the other for aspiration, were placed along the edges of the Transwell^®^ insert for perfusing oxygenated artificial cerebrospinal fluid (ACSF) 5 over tissue samples. For recording compound action potentials (CAP), a pulled glass micropipette electrode (1-4 MΩ) was inserted into the bulb of the channel near the clustered cell bodies, and the axons growing in the channel were stimulated with a concentric bipolar platinum-iridium electrode positioned 1-3 mm distal to the bulb. A platinum recording electrode was placed in the ACSF-filled glass micropipette and was connected to an amplifier set at 100x gain and 0.1Hz high pass to 3kHz low pass filtering. Stimulation pulse height and width were kept at 10 volts and 200 μsec, respectively. Samples were stimulated at a maximum repeat rate of 1 Hz, and at least 50 stimulations were applied per sample. Using an analog-to-digital converter (PowerLab; AD Instruments, Colorado Springs, CO), CAP waveforms were visualized and further stored using LabChart software (AD instruments). After CAP recording, using a stereomicroscope and camera, a snapshot of the stimulating and recording electrodes were taken to determine the distance between them for nerve conduction velocity (NCV) calculation. Latency was measured by subtracting the location of stimulus artifact by CAP peak location. NCV for myelinated hMN/hSC co-cultures and unmyelinated hMN monocultures was evaluated by dividing the distance between stimulating and recording electrodes by latency.

### Statistics

One-way analysis of variance (ANOVA) with Tukey *post-hoc* test was conducted using GraphPad prism software (GraphPad Software, Inc., La Jolla, CA, USA) to evaluate size differences between different kind of spheroids. For analysis of electrophysiology, mean and standard deviation were calculated and an unpaired *t* test was performed (GraphPad Software.) A p-value ≤ 0.05 was used to assign significant differences between the means.

## Results

### Schwann cells enhanced assembly of neurons into spheroids

In order to create a spheroid which fits appropriately within the dimensions of the Nerve-on-a-chip (NoaC) system (i.e., less than 1,000 μM in diameter, and maintain a high number of cells), we fabricated spheroids with different cellular densities. We also compared the sizes of different spheroids to understand the interactions between the hNs and hSCs.

After placing the desired number of cells in low attachment, round-bottom plates, we monitored the formation of spheroids every day. Mono-cultures of hSCs formed spheroids within approximately 2 days and were found to be regular in shape with sharp edges (Fig. 2a-2c). In contrast, mono-cultures of hNs did not self-assemble into spheroids in 2 days and instead formed many smaller spherical structures (Fig. 2g-2i). In co-cultures, hSCs facilitated incorporation of hNs into spheroids (~2 days) when compared to spheroids formed from hNs alone (~3-9 days, Figure 3). The edges of the co-culture spheroids (Fig. 2d-2f) were less sharply defined compared to spheroids containing the hSCs alone possibly due to the non-homogeneous nature of the coculture spheroid. Interestingly, the size of co-culture spheroids comprised of 75,000 hNs and 75,000 hSCs (1025±52 μm) was found to be very similar to the size of 75,000 hSCs alone (967±51 μm), suggesting that co-culture spheroids were more tightly packed and thereby confirming that the two cell types had affinity for each other.

**Figure 2.**
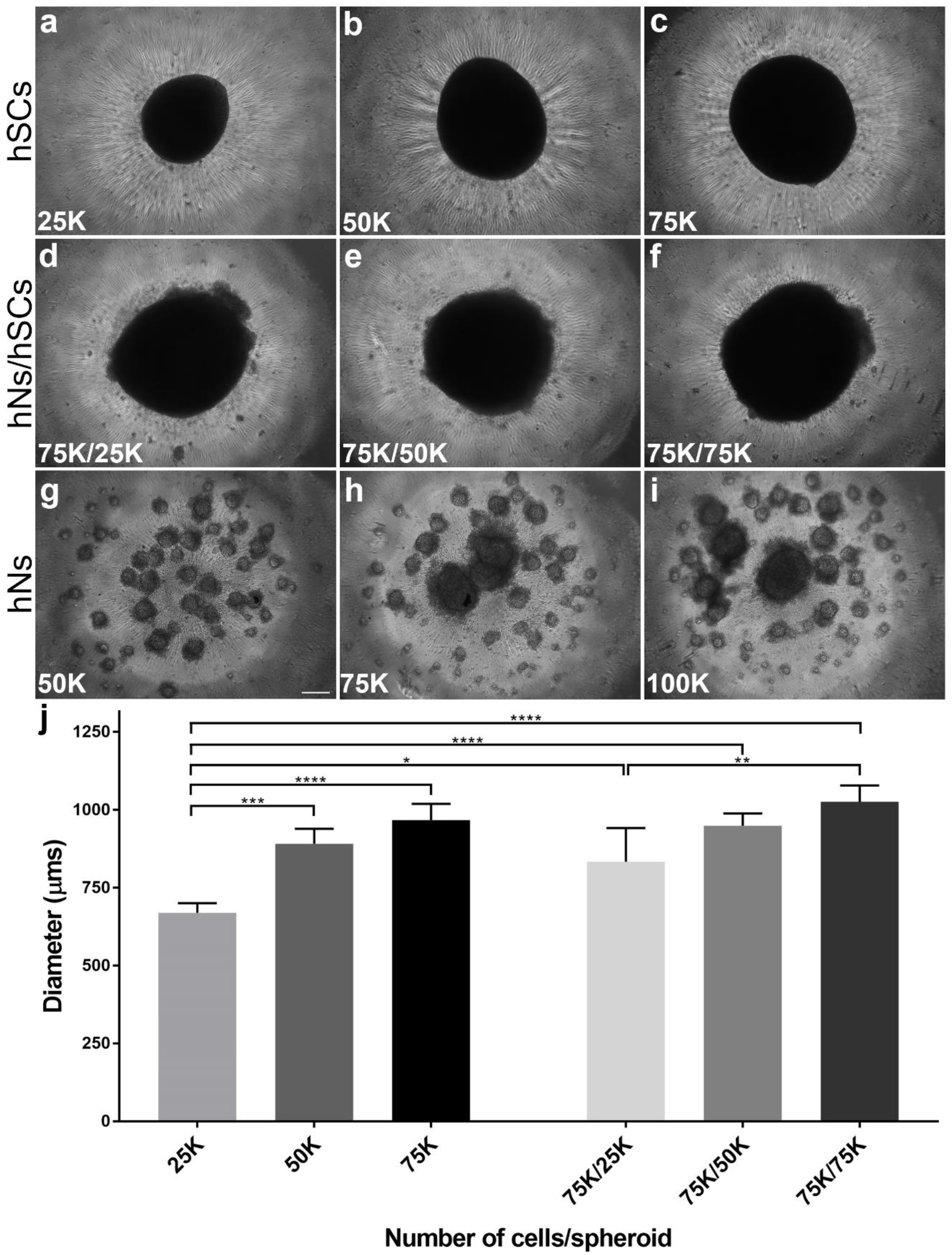
Fabrication of spheroids consisting of human neurons (hNs) and/or human Schwann cells (hSCs) after 2 days *in vitro*. Presence of hSCs expedited spheroid formation in the co-culture system while the hN-only condition did not form spheroids in 2 days. hSC spheroids were fabricated with three different cellular densities: 25,000 (a), 50,000 (b) and 75,000 (c). Co-culture spheroids were created with a constant hN density (75,000) but a changing hSC density: 25,000 (d), 50,000 (e), and 75,000 (f). In parallel, hN spheroids were fabricated at three different densities: 50,000 (g), 75,000 (h) and 100,000 (i). (j) Comparison of diameters of different spheroids revealed that co-culture spheroids were more compact as compared to mono-culture spheroids, showing that the affinity of hSCs for hNs resulted in a more tightly packed cluster of cells. K (multiple of thousand). N=4 and error bars represent standard error of the mean (SEM). Scale bar: 100μm. **** (p-value ≤ 0.0001), *** (p-value ≤ 0.001), ** (p-value ≤ 0.01), * (p-value ≤ 0.05).

By measuring the diameters of each of the different spheroid types (Fig. 2 and Fig. 3), we determined that having 75,000 neurons is the optimal number of hNs for constituting the hNoaC system as the spheroid size was found to be about 833±108, 948±39 and 1025±52 μm when we created co-culture spheroids with 25,000, 50,000 and 75,000 hSCs respectively. Neuron-only cultures revealed that spheroids increased in size predictably as the number of cells increased. All four hNs conditions (Fig. 3) saw a successive, significant increase in the spheroid size revealing that packing density across the four spheroids (25K, 50K, 75K and 100K) does not vary substantially and that the total number of cells contributes to spheroid size more than the interaction between the various cell types in the spheroid.

**Figure 3.**
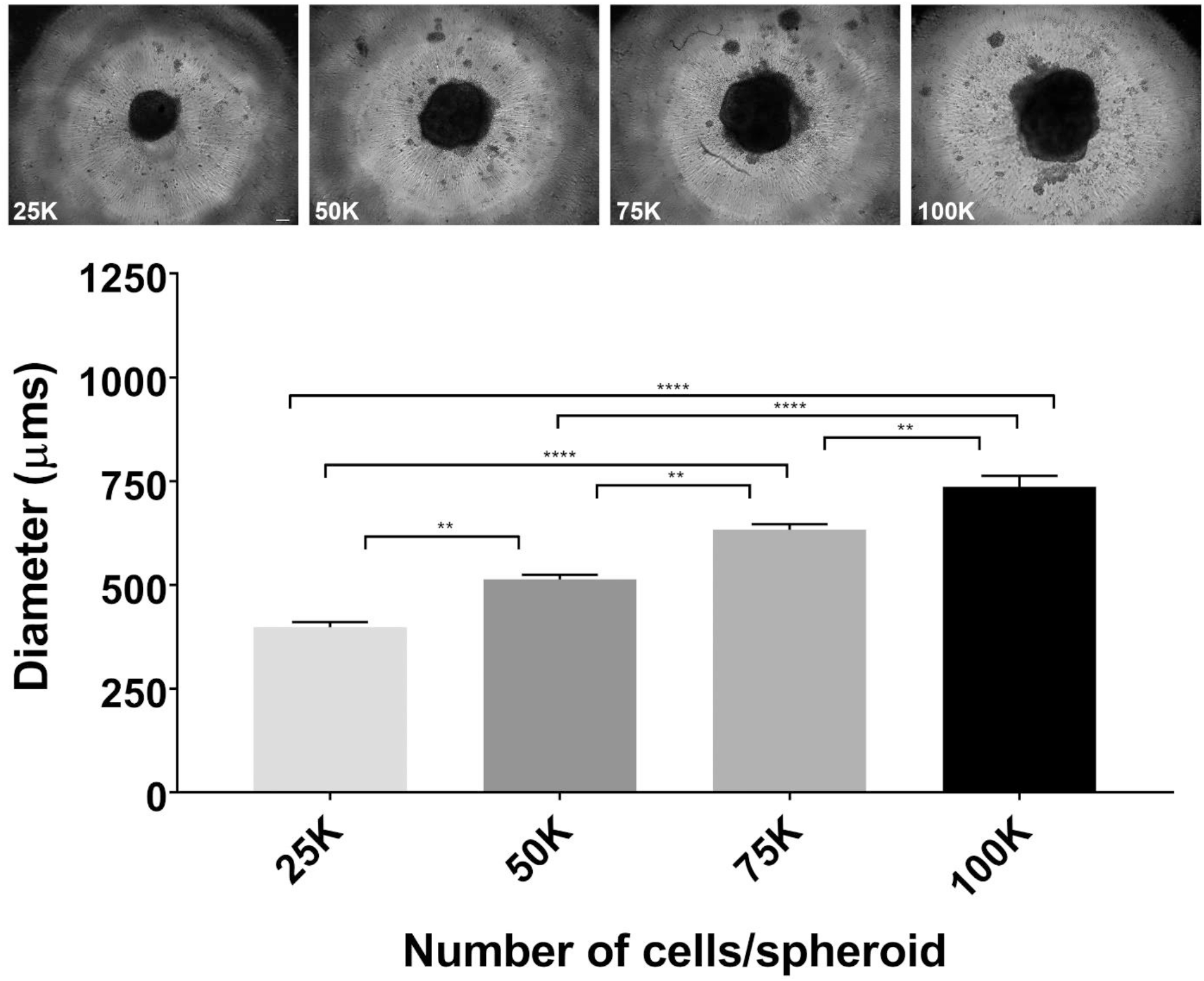
Fabrication of spheroids consisting of human neurons (hNs) alone. Qualitative inspection at >2 days after initiating the hN-only culture showed a consistent significant increase in spheroid sizes as the total number of cells increase (25K, 50K, 75K and 100K). Graph comparing the individual spheroid diameters supported the qualitative inspection. K (multiple of thousand). N=4 and error bars represent standard error of the mean (SEM). Scale bar: 100μm. **** (p-value ≤ 0.0001), ** (p-value ≤ 0.01).

### Co-culture spheroids showed robust neurite outgrowth in the NoaC system

While the outer portion of the dual hydrogel system was constructed with growth-resistant 10% PEGDMA, the inner part of the channel was filled with fully concentrated (8-12 mg/mL) Matrigel as a growth-promoting substrate. After gel formation, spheroids were gently transferred on top of the bulb part of the channel and left to grow in medium containing 10% FBS but lacking NGF in order to enhance the proliferation and migration of hSCs while delaying neurite extension from hNs. After a week, the incubation solution was switched to medium supplemented with NGF and L-ascorbic acid to facilitate neurite growth and myelination by the hSCs in contact with the growing axons.

Confocal imaging revealed the 3D nature of the reconstituted *in vitro* nerves and showed that both cell bodies and axons were present throughout the depth of the channel (Fig. 4). Neurites grew an average of ~1 mm every week (Supplemental fig. 1). Addition of FBS was a key factor in optimizing myelination as the basal media with ascorbic acid but without FBS did not support hSC migration and myelination (data not shown). Immunostaining after four weeks with S100 revealed that hSCs cells migrated about 1-1.5mm outside the spheroid and elongated along the growing axons (Fig. 4A-C), while the axons reached to the very end of the Matrigel-filled channel (~5mm). Interestingly, for many co-culture samples, spheroids influenced the growth of axons such that they appeared to turn back after growing for some distance, possibly because of the chemotactic effect created by growth factors released from hSCs in the spheroid (Supplemental fig. 2). The effect on axons was more prominent as the number of hSCs in the spheroid increased.

**Figure 4.**
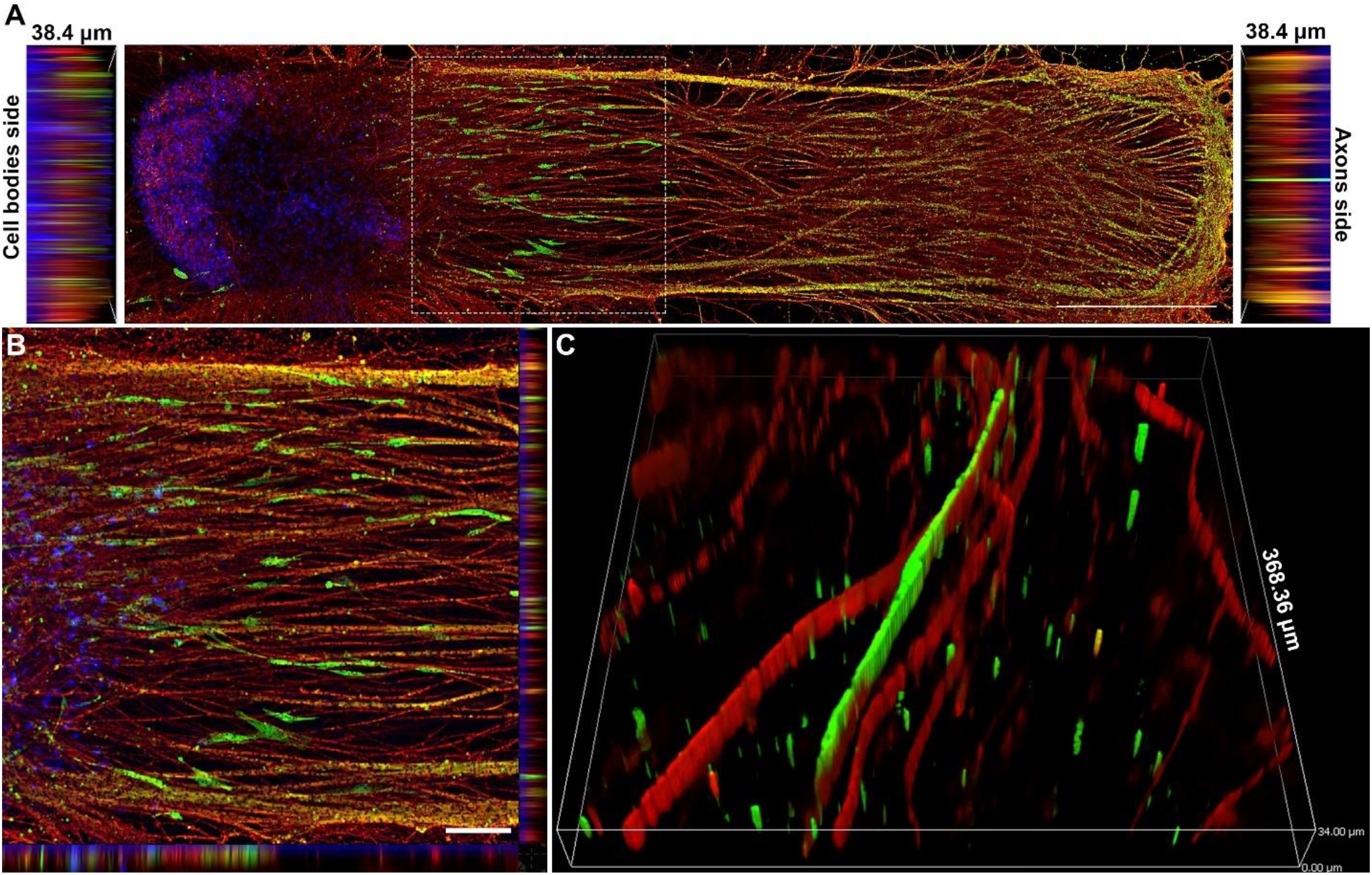
Schwann cells migrated out of the spheroid and elongated along the axons. (A) Image showing how human Schwann cells (hSCs) stained for the hSC marker S100 (green) migrated out of the spheroid along with growing axons stained for βIII-tubulin (red) over a period of 4 weeks. Nuclei were labeled with DAPI (blue). Scale bar: 1000 μm. (B) High-magnification image of inset from image A. Scale bar: 25 μm. (C) 3D image showing close-up of the relationship between hSCs (green) and myelinated axons (red). Slice size was 368.36 × 368.36 × 34.00 μm.

### Myelination and Nerve Fiber Structure of *in vitro* Human Nerves

Finally, along with immunostaining and confocal microscopy, we also performed plastic resin embedding and sectioning to evaluate the level and quality of myelination in the system with TEM. Evidence of effective myelination in the system included but was not limited to non-compacted myelin (Fig. 5A), compact myelin (Fig. 5B), and myelin in the process of compacting (Fig. 5C). For the axons, where we saw evidence of myelination, the G-ratio of myelinated nerve fibers was 0.57±0.16. Axonal diameters of the myelinated and unmyelinated axons were 0.55±0.33 and 0.40±0.15 μm, respectively. We also saw evidence of laminar myelin formation without axons (Fig. 5D), the presence of intracytoplasmic lamellar bodies (Fig. 5E), and naked (unmyelinated) axons (Fig. 5F). The appearance of the intracytoplasmic lamellar bodies, which consisted of membrane whorls with relatively regular spacing, was interpreted to represent autophagosome production consistent with recycling of senescent organelles. The distribution of lamellar bodies was sparse and apoptotic nuclei were not observed, indicating that affected cells were not engaged in programmed cell death.

**Figure 5.**
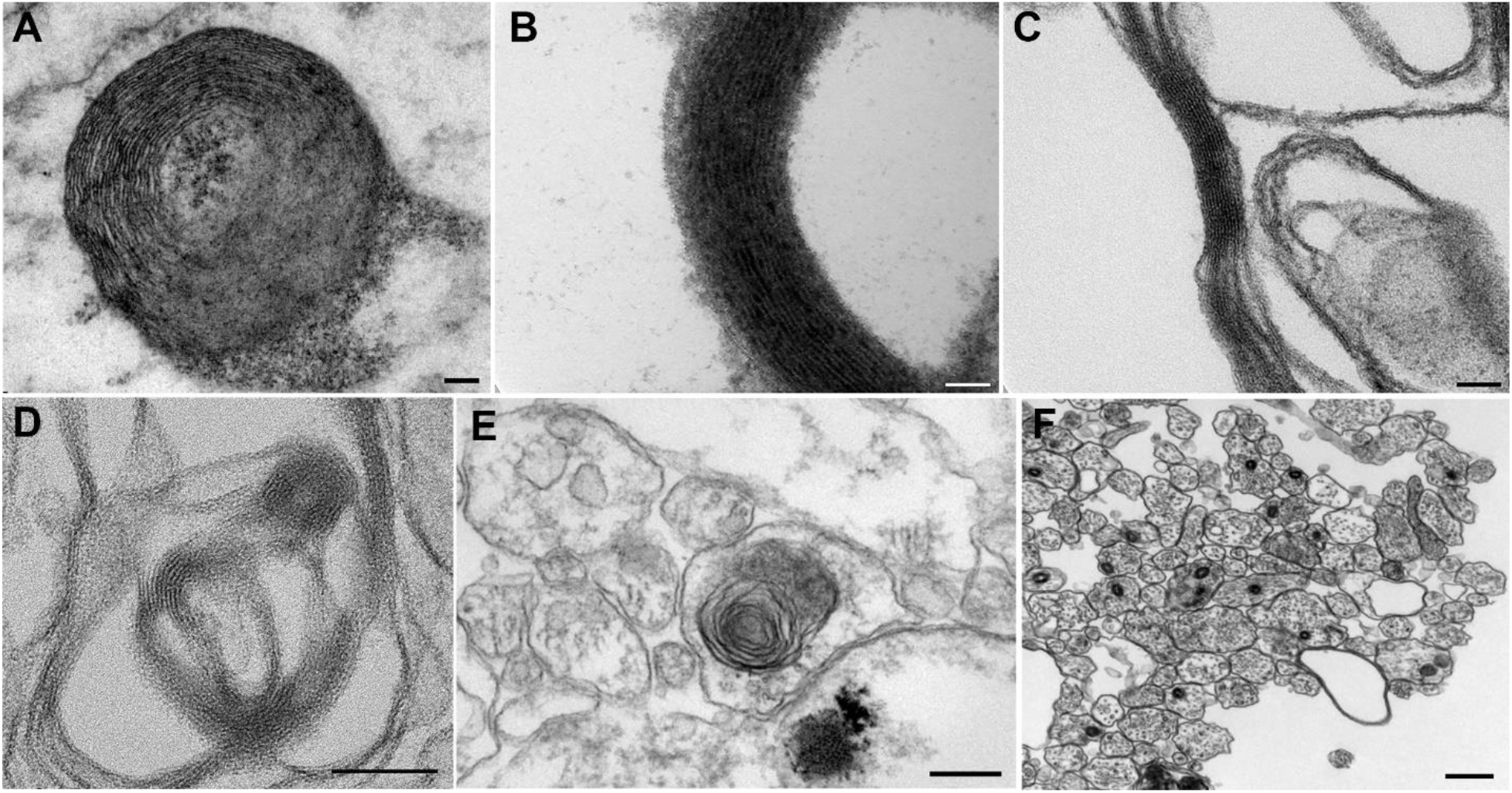
Various stages of myelin formation observed in *in vitro* reconstituted human nerve. (A) Non-compact myelin. (B) Compact myelin. (C) Myelin in the process of compaction. (D) Myelin formation without axons. (E) Intracytoplasmic lamellar bodies. (F) Naked (unmyelinated) axons.

### *In vitro* human nerves exhibit effective, composition-dependent electrical conductivity

To determine whether we can measure nerve conduction velocity (NCV) of iPSC-derived human neurons (hNs) with or without human Schwann cells (hSCs), we used a technique similar to brain slice electrophysiology. We stimulated the axons inside the channel and recorded the compound action potential (CAP) from the cell bodies (Fig. 6A). Axons were stimulated about ~1-3 mm away from the cell bodies, and the distance the impulse traveled was calculated between the stimulating and recording electrodes. We evaluated two types of NCV, onset NCV and peak NCV, in order to determine the difference between the fastest signal and the peak signal (Fig. 6B’-6B’’). To our surprise, we found the onset and peak NCV with the hN/hSC co-culture samples was slower compared to hN mono-culture samples. Onset NCVs for 75K hNs mono-cultures and 75K/25K hNs/hSCs co-cultures were determined to be 0.28±0.07 and 0.20±0.02 m/s, respectively, while the peak NCVs were found to be 0.18±0.04 and 0.13±0.02 m/s, respectively (Fig. 3C). Onset and peak NCVs were determined with difficulty from samples where the number of SCs were higher (co-cultures of hNs/SCs at 75K/75K and 75K/50K). Qualitative inspection of the samples revealed slightly less dense neurite outgrowth with co-culture samples, which can reduce the NCV.

**Figure 6.**
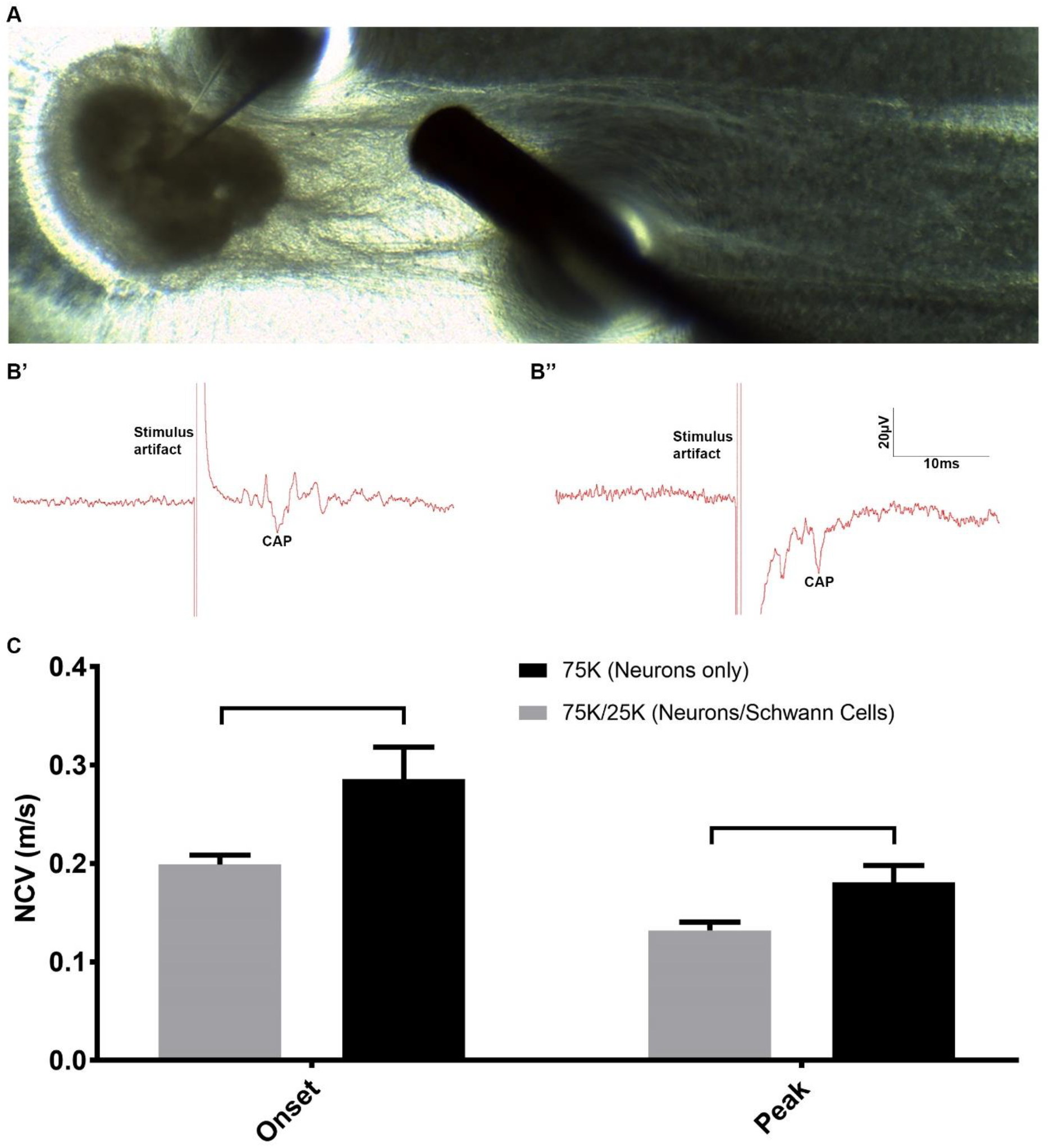
Electrophysiological testing of human nerve-on-a-chip (HNoaC). (A) Image showing the placement of stimulating and recording electrodes for recording compound action potentials (CAP) from *in vitro* HNoaC samples. (B’) Electrical trace showing the CAP generated in spheroid containing only 75,000 human neurons (hNs). (B’’) Trace showing the CAP generated in a co-culture spheroid consisting of 75,000 hNs and 25,000 human Schwann cells (hSCs). (C) Graph showing comparison of Onset and Peak nerve conduction velocities (NCV) between hNs-only monoculture (75K neurons) and co-culture (75K hNs with 25K hSCs). Onset NCV shows the start of the onset potential, while the Peak NCV shows the NCV for the biggest action potential. Nerve conduction velocity was observed to be slower in co-culture spheroids as compared to hNs-only spheroids (p-value ≤ 0.05). Error bars represent standard deviation (SD).

## Discussion

In this study, we present the first biomimetic, all-human *in vitro* model of peripheral nerve, assembled as a Nerve-on-a-Chip (NoaC) platform. This microengineered dual hydrogel system retains the neuronal cell bodies in a defined location (i.e, the “ganglion”) and confines dense 3D axonal outgrowth within a narrow channel that extends linearly (i.e., the “nerve”) away from the clustered cell bodies. The system supports current “gold standard” functional (e.g., electrophysiological testing) and structural (e.g., qualitative and quantitative microscopic analyses) endpoints required for assessing neuropathological conditions associated with peripheral neuropathies, which represent a growing medical concern. Innovative aspects of this study include reproducible fabrication of neuron-Schwann cell co-culture spheroids, robust viability (~4 weeks) and extensive neurite outgrowth (~5 mm) *in vitro*, effective myelination of human iPSC-derived neurons (hNs) by primary human Schwann cells (hSCs), and the ability to measure nerve conduction velocity (NCV) in an *in vitro* setting amenable to human disease modeling, drug discovery, and toxicity screening.

### Challenges in Producing In Vitro Nerve Systems

*In vitro* myelination using primary hSCs has long been a challenge, due in part to complications associated with extracting hSCs from adult nerves ^8,9^, contamination by fibroblasts ^8-10^ and the transformation of SCs to a proliferative/non-myelinating phenotype *in vitro* ^11,12^. Co-culture conditions are well established for myelination of rat dorsal root ganglion (DRG) sensory neurons by embryonic, neonatal, and adult rodent SCs ^13-15^. However, similar co-culture conditions fail to recapitulate myelination using human SCs cultured with rat DRG neurons ^11^. Rigorous purification of primary human SCs or differentiation of human stem cells or human fibroblasts to SC-like cells results in limited levels of myelination of rat sensory neurons, but the extent seen in the mixed-species cultures is significantly less compared to that achieved using embryonic rat SCs ^11,16^, possibly due to species differences or density of SCs compared to the number of axons. Recently, Clark and coworkers ^17^ successfully demonstrated myelination of human stem cell-derived sensory neurons by rat SCs. Still, an *in vitro* system exhibiting myelination of human iPSC-derived neurons by human Schwann Cells has remained elusive.

In the last few years, many studies focused on creating neuronal-glial organoids to create brain-like tissue *in vitro^18-22^*. Interestingly, all these strategies focused on differentiating the aggregates of neural progenitor cells into more defined neural structures. In contrast, we reverse-engineered the process by bringing together two differentiated cell types to evaluate their interactions with each other and the potential of self-assembly. To mimic the growth of embryonic dorsal root ganglia (DRGs) *in vitro*, we produced neuron-Schwann cell spheroids using ultra-low attachment, 96-well plates to facilitate crosstalk between axons and SCs, which is important for the differentiation of SCs toward a myelinating phenotype ^23,24^; thus, by bringing axons and SCs close together in a 3D spheroid, we enhanced the chances of cross communication and successful myelination. Following addition of an anti-oxidant, ascorbic acid, we observed the first-ever evidence of myelination *in vitro* of stem cell-derived human neurons by primary human Schwann cells. Both hNs and hSCs had different rates of self-assembly and spheroid fabrication individually, but when put together hSCs enhanced the quality and improved the speed of spheroid self-assembly as compared to the neuron-only condition. Based on spheroid diameter, coculture spheroids were found to be more compact as compared to either hNs or hSCs spheroids showing the enhanced interaction between the two cells types.

### Migration of Schwann cells out of the spheroids

Schwann cell migration is a critical phenomenon during development and peripheral nerve regeneration following injuries ^25^. Cues that direct the fate of neural crest cells to Schwann cell precursors and ultimately to Schwann cells are largely unknown; however, it has been known for decades that both precursor cells and Schwann cells rely on growing axons for differentiation, proliferation and functional maturation ^26^. Here, for the first time, we were able to observe this migration *in vitro* for tissues of human origin by creating this mini-ganglion comprised of hNs and hSCs. Axons extended outwards along with migrating hSCs which aligned themselves with the growing axons in this process. It was interesting to observe that hSCs only migrated to about ~1 mm outside the spheroid as compared to total axonal growth of about 5 mm. This could possibly be due to a lower number of hSCs relative to neurons added during these experiments as compared to typical 2D co-culture experiments, where the usual convention is to add >100,000 Schwann cells in a smaller 2D area ^17,27^. This modest extension of hSC compared to axon outgrowth could also be a result of only one week of pre-myelination period and addition of NGF to the media after the first week of growth. NGF has been shown to enhance neuron-Schwann cell interaction and also myelination ^28^, and thus could be a factor which reduced the migration of SCs. Based on the migration of human SCs outside the spheroid, we hypothesize that this HNoaC model can also be used for studying the migration potential of SCs in the presence of therapeutic molecules and thus create possible therapies for patients with peripheral nerve injuries.

### Mini-Nerve Structure

hSCs are known to behave differently *in vitro* as compared to rat Schwann cells in terms of their reactivity to mitogens and growth factors as well as their failure to recapitulate myelination ^29^. Our 3D spheroid model of human nerve exhibited typical features of nerve trunks observed in nerves acquired during autopsy or biopsy procedures. Axons had a complete complement of organelles including cytoskeletal filaments and mitochondria, and often but not always were associated with sheaths of myelin characterized by closely approximated myelin laminae. The apposition of myelin layers varied among nerve fibers, and in some cases laminar myelin formed in the absence of axons; both these findings are rarely encountered in differentiated nerves harvested *in vivo*, indicating that some differences in differentiation state do occur (as expected) in culture. That said, sufficient numbers of myelinated axons were observed in the mini-nerve portion of the co-culture system to render it a suitable surrogate for mixed somatic nerves (i.e., those containing densely myelinated, thinly myelinated, and unmyelinated axons).

### Evaluation of nerve conduction velocity (NCV)

Different neuropathies are known for showing different kinds of neurophysiological characteristics ^30^. *In vitro* microengineered nerve thus should be capable of defining electrophysiological changes as a way of conducting investigative and mechanistic toxicological studies. In this study, which is the first to use human iPSC-derived neurons to study nerve conduction, we were able to see differences in the nerve conduction velocity (NCV) between the myelinated and unmyelinated human axons which shows that this system is sensitive enough to evaluate nerve function. To our surprise, myelinated hNs/hSCs co-culture samples showed a slower NCV as compared to unmyelinated hNs-only mono-culture samples. A qualitative inspection of the culture revealed that the number of hSCs in the spheroid may have reduced the outgrowth and axonal density in the channel of HNoaC, which could easily result in reduction of NCV. Also, because many axons appeared to be turning back in high SCs (50K and 75K, Supplemental figure 2) density co-cultures, we were not able to determine the most optimal length between the point of stimulation and point of recording which can impact the NCV calculations. Furthermore, the presence of non-neuronal cell bodies in co-culture spheroids decreases the probability of recording from the appropriately stimulated cell bodies. Importantly, NCV from hNs was found to be considerably lower as compared to the NCV values obtained in human patients^31-33^. This is not particularly surprising, considering an *in vitro* system, at room temperature, comprised of iPSC-derived neurons that are less mature as compared to mature, myelinated axons of nerves assessed *in vivo*.

The simple design of this fully Human NoaC system opens new avenues in translational research. The platform can be used not only for screening drug candidates on the basis of clinically relevant electrophysiological and histopathological metrics but can also be used for investigating basic mechanisms driving nerve diseases, including but not limited to toxic, demyelinating, and other neurodegenerative conditions. For a given manipulation or treatment, comparison of data acquired from our conceptually identical Rat NoaC^6^ and Human NoaC will help close the gap between nonclinical testing and our ability to anticipate responses and potential safety risks in humans.

## Acknowledgements

We would like to acknowledge Ying Xiao from Louisiana State University for her help on histology and transmission electron microscopy imaging. Funding for this research was provided by grants from the NIH (R42TR001270) and CASIS (GA-2016-238).

## Author contributions

ADS, LM, EJ, HW and JQB conducted the experiments. ADS and EJ analyzed the data and prepared the figure plates. ADS, HN, JLC and MJM designed various parts of the study. BB advised on the nerve structure. ADS, LM, EJ, BB, JLC and MJM wrote various sections of the manuscript. All authors reviewed the manuscript.

## Competing interests

The author(s) declare no competing interests. JLC and MJM are co-founders of AxoSim, Inc. with equity stakes in the company. ADS, LM, EJ, HW, JQB, and HN are paid employees of the company. BB is a consulting pathologist.

## Data availability

The datasets generated during and/or analyzed during the current study are available from the corresponding author on reasonable request.

